# Episodic memories: how do the hippocampus and the entorhinal ring attractors cooperate to create them?

**DOI:** 10.1101/2020.04.25.061614

**Authors:** Krisztián A. Kovács

## Abstract

The brain is capable of registering a constellation of events, encountered only once, as an episodic memory that can last for a lifetime. As evidenced by the clinical case of the patient HM, memories preserving their episodic nature still depend on the hippocampal formation, several years after being created, while semantic memories are thought to reside in neocortical areas. The neurobiological substrate of one-time learning and life-long storing in the brain, that must exist at the cellular and circuit level, is still undiscovered. The breakthrough is delayed by the fact that studies jointly investigating the rodent hippocampus and entorhinal cortex are mostly targeted at understanding the spatial aspect of learning. Here we present the concept of an entorhinal cortical module, termed EPISODE module, that could explain how the representations of different elements constituting episodic memories can be linked together. The new model that we propose here reconciles the structural and functional observations made in the entorhinal cortex and explains how the downstream hippocampal processing organizes the representations into meaningful sequences.

## 1. Introduction

As the clinical case of the patient HM has proven, the medial temporal lobe memory system, including the hippocampus proper and the adjacent entorhinal, perirhinal, and parahippocampal cortices, is critically concerned in the retention of new episodic memories (Scoville and Milner, 1957; Squire, 2009). The intuition dictates that the function of such a memory system must consist of linking different neural representations together and, also, at least in some cases, organizing them into a sequence.

It is hypothesized that the link established between representations in the medial temporal lobe system is projected back to neocortical areas over a longer period of time. Nonetheless, some authors proposed that some autobiographical memories depend on the medial temporal lobe system as long as they can be recalled (Steinvorth et al., 2005), a view that has been criticized later without questioning that the transfer to neocortical areas still can take a few years (Squire, 2009).

The circuit-level mechanism of creating a stable association in the medial temporal lobe system must involve Hebbian plasticity between neurons and is most likely based on spike timing dependent plasticity (STDP) that has been widely studied *in vitro* and in animal experiments. Although rules for STDP are not as straightforward to define in the hippocampus as are in the neocortex, it is still established that STDP can strengthen the connection between signals that are separated by no more than a few dozens of milliseconds (Nishiyama et al., 2000). One key question, therefore, is how the brain can forge different elements together into a single, coherent episodic memory when the temporal separation of such elements can be substantially larger than one minute. An intriguing observation, viewed by many scientists as a solution, is that the representations of temporally separated events are brought much closer to each other within the approximately 100 ms long theta cycle in rodents (Drieu and Zugaro, 2019). The so-called theta sequences not only bring the elements of an episodic memory closer in time but the fine temporal code expressed in one theta cycle can be repeated several times during an episode to be remembered (Drieu and Zugaro, 2019). Indeed, studies of the hippocampal STDP have shown that positive plasticity is not only dependent on how separated the two stimuli to be paired are, but also on how many times and at what frequency the pairing is repeated (Wittenberg and Wang, 2006). Therefore theta sequences appearing in wakeful periods are well-positioned to elicit hippocampal plasticity. However, on the other hand, the mechanism by which representations of events, places, items, stimuli are placed next to each other on a theta cycle is not completely understood. The prevailing view is that hippocampal theta phase precession (O’Keefe and Recce, 1993), now thought to originate from the entorhinal cortex (Schlesiger et al., 2015), is the most important phenomenon contributing to the emergence of hippocampal theta sequences (Jaramillo and Kempter, 2017). In the present article we put forward a model that rather views the CA1 theta sequences as a „by-product” of Hebbian association of the memory elements in the hippocampus and points to the phase precession as the primary mechanism that drives such association in the CA3.

The heart of the model is a unit circuit in the entorhinal cortex that we have baptized the EPISODE module (also an acronym of the previously identified cell types it is built from). The EPISODE module provides an explanation for the generation of phase precession, for the emergence of grid cells and for the theta cycle skipping (Brandon et al., 2013). Furthermore, the phase precession and the fine temporal code along the theta cycle produced by the EPISODE module can be translated in downstream hippocampal areas into representations that are optimally separated for efficient STDP that, in turn, makes heteroassociation and sequence generation possible as will be discussed briefly.

The unique feature of the new model is that it assumes the physical presence of 1-dimensional ring attractors built from reelin-positive cells in the entorhinal cortex. The emergence of 2-dimensional activity seen in grid cells is explained by combining more 1-dimensional units rather than arranging the neurons physically in torus topology. The model unifies some features of ring attractor models and some of the oscillatory interference models to explain the emergence of both the phase precession and of the hexagonal grid activity in a similar way to already published concepts (Navratilova et al., 2012). However, in our view, the general function of the 1-dimensional ring attractors extends far beyond spatial coding and is rather centered on assigning a theta phase in humans to any relevant event, place, item, place or stimulus that is to become a part of any episodic memory.

## 2. The entorhinal component of the episodic memory circuit: the EPISODE module

### 2.1. Rodent grid cells: what mechanism explains their activity and how are they anatomically organized?

The discovery of rodent grid cells (Hafting et al., 2005; Sargolini, 2006) let a technically relatively easy functional insight into a subset of a neuronal population that we propose to be of key importance for registering episodic memories. Grid cells show a strikingly regular spatial activity: when the animal navigates in an environment, the locations where these entorhinal cells become active are arranged in a hexagonal pattern (Hafting et al., 2005; Sargolini, 2006). Such activity proved to be hard to explain given the almost perfectly regular 2-dimensional nature of the hexagonal pattern which is unaffected by the speed of the animal (figure 1), and the omnidirectional theta phase precession in each firing field which is bimodal (Climer et al., 2013) for the neurons in the layer II of the medial entorhinal cortex (MECII). Two main classes of models were elaborated to elucidate grid cell activity: the oscillatory interference models and the continuous attractor neural network models. The first group hypothesizes the interaction of two different oscillatory signals, similarly to those generating the Doppler’s effect, one is supposed to be a signal modulated by the speed of the animal, in most of the models this is proposed to be the theta oscillation of the LFP, the other signal is the intrinsic resonance of the stellate cells in the MECII resulting from their biophysical properties. The main criticism these models have to face is that grid cells have also been found in the bat entorhinal cortex where regular theta oscillation is practically absent (Yartsev et al., 2011). A second strong general argument against the whole class of these models is the implausibility of maintaining oscillations of different frequencies in electrically compact cells such as the MECII stellate cells or any other known cell type of the entorhinal cortex (Remme et al., 2009). The other major class of gird cell models, the continuous attractor neural network models, assume an active center that is traveling in a population of neurons arranged in a structure (for example a torus) that permits propagation without boundary and each neuron within the structure would fire each time when the activity circling in the network travels through it (McNaughton et al., 2006). Neighboring neurons are supposed to code for adjacent values of a parameter (such as head direction or the spatial position in the case of grid cells). These models on the other hand, as detailed below, require an exquisite wiring of the neurons (including those within the 2-dimensional structure itself and those that drive the center of activity forward), unprecedented elsewhere in the mammalian central nervous system. It is especially hard to imagine that such extraordinary wiring can form at the state of the ontogenesis when the adult-like grid cell activity develops abruptly (Wills et al., 2012).

**Figure 1.**
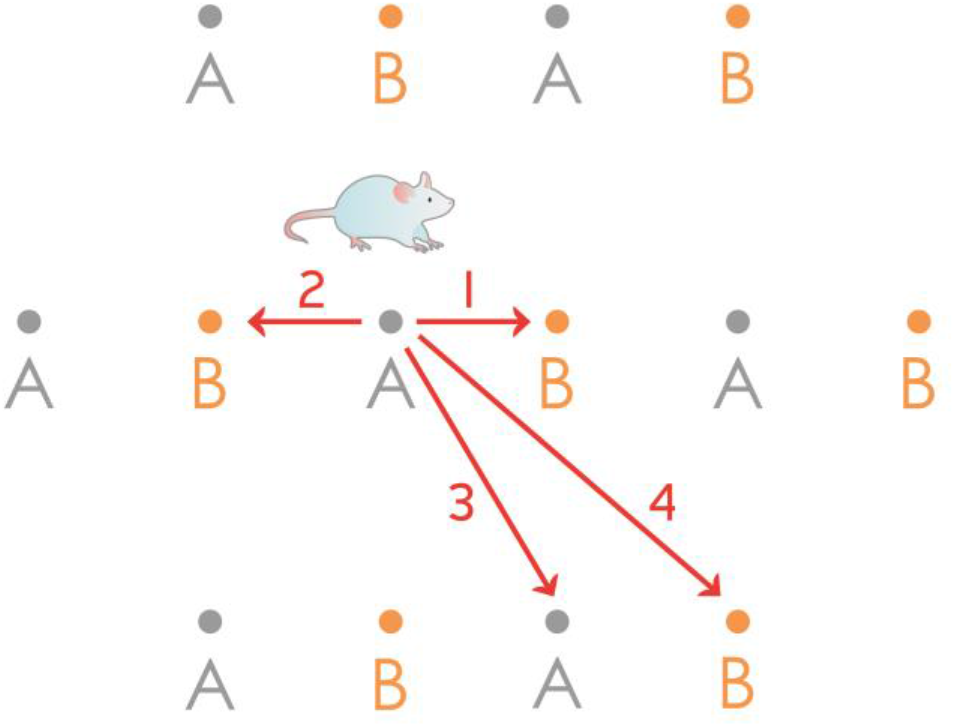
Rodent grid cell activity cannot be explained by a simple 1-dimensional ring attractor. The figure shows the firing fields of two grid cells („A” and „B”) from the same hypothetical 1-dimensional ring attractor, having equal grid scales but different spatial phases. Forward and backward movement along a linear track could easily be interpreted: when the animal starts its journey from the center of the field of the neuron „A”, the bump can travel the half of the distance of the whole attractor either clockwise or anticlockwise (routes 1 and 2) and reach the same neuron „B”. However, trajectories along 3 and 4 cannot be reconciled with this 1-dimensional model: 3 would guide back the bump to the same neuron („A”) while 4, a slightly longer route would guide the bump only halfway around the ring attractor to reach the neuron „B”.

It is not only the firing rate map of the grid cells that is strikingly regular but the layer II of the medial entorhinal cortex where the neurons are physically located also shows regular, periodic cytoarchitecture, best seen in tangential sections. This prompted several authors to find the correlation between anatomical and electrophysiological regularity (Naumann et al., 2018; Ray et al., 2014). Early on, an isomorphic mapping hypothesis has been published (Brecht et al., 2013) that was pioneering in the sense that it directly correlated the spreading of the neuronal activity in an anatomically well-circumscribed MECII network and the pattern of activity seen in real space when the animal navigates. However, several contradictions can be identified in this particular concept, most of them arising from results that subsequent experiments produced. The most important of those include (a) the identity of the grid cells, (b) the assumed connectivity, (c) the organizational level of regularity. The hypothesis (Brecht et al., 2013) identified grid cells as the calbindin positive pyramidal cells of the MECII, later also termed „island cells” (Kitamura and Tonegawa, 2014; Sun et al., 2015). These neurons are grouped into discrete patches that are arranged into a network that is visible in tangential brain sections and its structure is close to hexagonal in many species. In contrast, a recent investigation that directly observed functioning grid cells using calcium imaging (Gu et al., 2018) clearly concluded that most grid cells were reelin positive stellate cells (75,6%), the other prominent cell type of the MECII, also termed ocean cells (Kitamura et al., 2015). The connectivity required for the isomorphic mapping hypothesis was also unrealistically sophisticated: an isoposition connectivity was assumed between patches of the same spatial scale and across hemispheres so as to synchronize the positions of the centers of activity between patches. Moreover, „wrap-around” connectivity was also assumed within every single 150 μm large patch so that the attractor dynamics can be continuous and cover a physical space of any size (Brecht et al., 2013). Although the topology of the „wrap around” connectivity was not directly specified, it must have been toroidal so that a truly 2-dimensional continuous attractor network can be built up within each calbindin positive patch. Finally the model is challenged by the fact that hexagonal layout of the calbindin positive neurons, the connectivity of which is assumed to map the physical space, has never been observed directly within a single patch. The anatomical hexagonality rather appears in the way the patches themselves are organized within the continuum of clean cells.

To reconcile the anatomical organization and the computational need that the observed grid activity implies we propose a 1-dimensional ring attractor in our new model that consists of interconnected MECII stellate cells arranged in a ring topology. These rings of neurons could encircle calbindin positive pyramidal patches, however, this is not an absolute requirement for the model to describe their activity. Indeed, recent direct observation of grid cell function (Gu et al., 2018) supports the idea that the actual rings can also be embedded in a field of other stellate cells, and their actual shape can very well differ from a regular circle. We believe that the fact that grid cell pairs show a correlation between their spatial phase of activity and their physical distance in the brain (Gu et al., 2018) underscores the idea that ring attractors of stellate cells could explain the grid cell firing patterns.

A prominent early model of grid cell firing described the continuous attractor network in a 1-dimensional scheme with mentioning the obvious extension to 2-dimensional variants (Navratilova et al., 2012). The appealing feature of this model is that it incorporates the intrinsic resonance and postspike dynamics of the MECII stellate cells, similarly to oscillatory interference models. Furthermore, the same model explains the generation of phase precession which is experimentally shown to arise in the MECII independently from hippocampal activity (Hafting et al., 2008) and to be inherited by the hippocampus proper (Schlesiger et al., 2015). Therefore our conviction is that the model by Navratilova et al. is the best description so far of the entorhinal grid cells and before extending it to general neuronal module subserving episodic memory, we briefly describe its features in the following section.

The concept and the *in silico* simulation presented by Navratilova et al. (2012) is built on a 1-dimensional ring attractor, however, it also incorporates the intrinsic resonance and other physiologically realistic properties of MECII stellate cells, thereby it explains not only the grid-like firing activity but also the emergence of phase precession. Originally it simulated 100 MECII stellate cells („grid cells”) arranged in a ring attractor and capable of exciting each other. The activity center („bump”) in the attractor was moved in one direction by 100 „north conjunctive cells” and in the other direction by 100 „south conjunctive cells”, these neurons received excitatory input from the grid cells to track the actual position of the animal. Further excitatory input reached the conjunctive cells from their congruent („north” and „south”) head direction cells. The asymmetric excitatory connections fed back to the grid cells moved the bump forward or backward in the ring of grid cells (figure 2). Theta oscillation was simulated as an 8Hz sine wave input onto the north and south conjunctive cells and global inhibition restricted the activity in the whole network. Importantly the medium afterhyperpolarization (mAHP, peaking at 20 ms after each spike) and afterdepolarization (ADP, peaking at 100 ms after each spike) of the grid cells were incorporated in the model. The simulations produced remarkable similarities with the *in vivo* observable properties of the network. First, within a fourfold range of head direction input strength, the speed of the bump within the attractor increased linearly when the voltage of the head direction input was increased, this result corresponds to the *in vivo* maintenance of the grid pattern regularity for faster and slower epochs of running. Second, crucially, phase precession that is shown to be generated in MECII experimentally (Hafting et al., 2008; Schlesiger et al., 2015), naturally arose in the simulations, and its observed omnidirectional nature (Skaggs et al., 1996) could be explained. Finally the ADP, as defined on the basis of the real, physiological properties of the stellate cells, resulted in the re-generation of the bump in each theta cycle, even in the absence of excitatory input from conjunctive cells. The fact that ADP and conjunctive cell input both contributed to the firing of grid cells could explain the bimodality of the phase precession (Yamaguchi et al., 2002).

**Figure 2.**
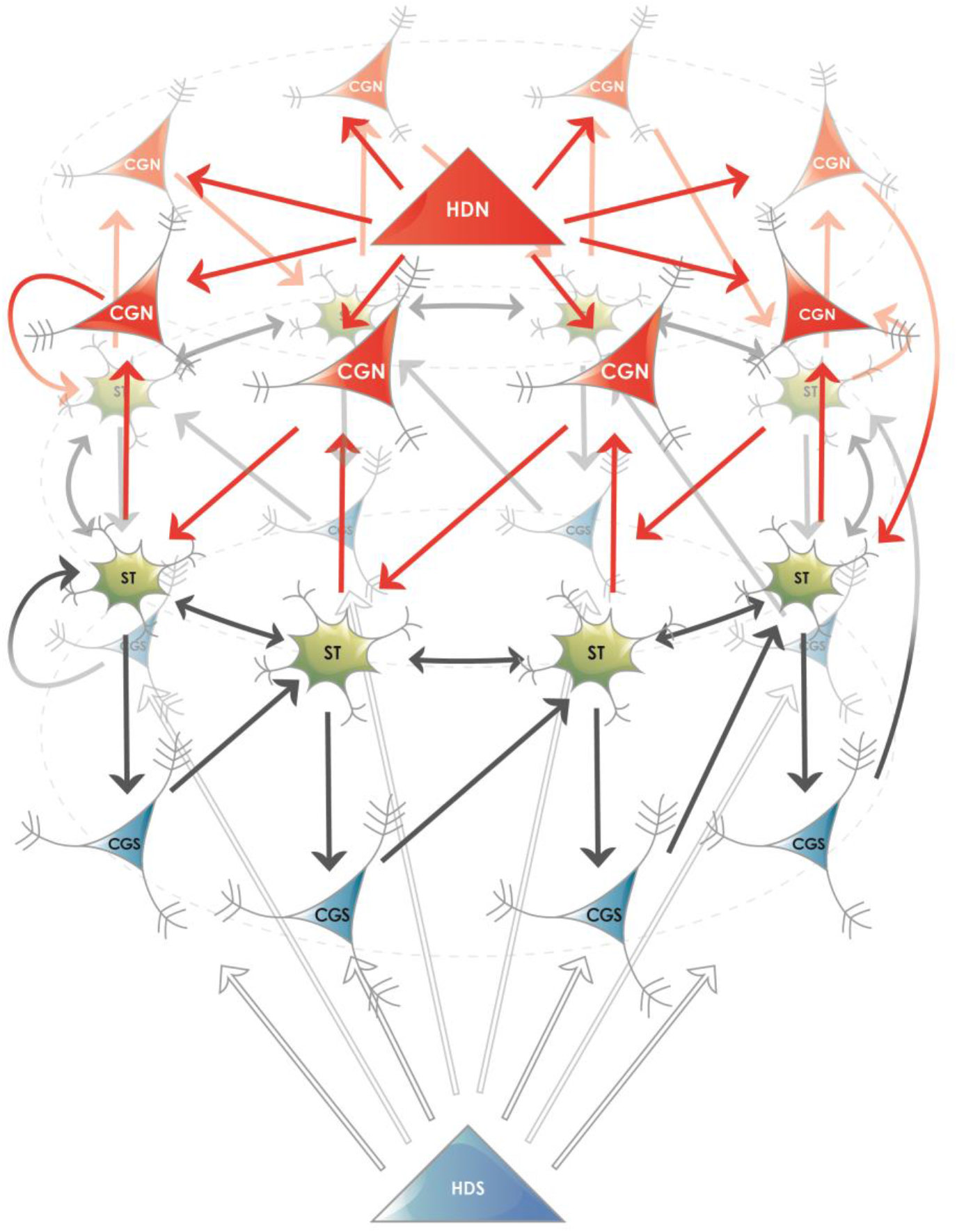
Proposed ring attractor in the superficial layers of the entorhinal cortex. Five different cell types are shown, their names are abbreviated according to the nomenclature in the MEC: (i) Stellate or fan neurons (<u>**ST**</u>), both named SODeR cells in our generalized model, a fraction of which are grid cells in rodents. (ii) Putative pyramidal cells from layer III (<u>**CGN**</u> and <u>**CGS**</u>), named PiPhA cells in our generalized model, capable of displacing the activity center in the ring of ST cells. A fraction of these neurons show conjunctive grid („CG”) cell activity in rodents, the north („N”) and south („S”) subtypes move the bump of activity in opposite directions. (iii) Superficial entorhinal neurons (<u>**HDN**</u> and <u>**HDS**</u>), both named ErReD cells in our generalized model, that provide continuous input to the CGN and CGS cells when an event is ongoing. Medial entorhinal head direction („HD”) cells are typical neurons falling into this category. For the sake of clarity several features are simplified in the figure: (i) the physical shape of the attractor is not supposed to be circular, the ring only displays its topology, (ii) the two antagonistic set of PiPhA cells (CGN and CGS in the depicted example) do not have to be located on the opposite sides of the SODeR cells located in layer II, actually their most likely position is below them (layer III) for both PiPhA cell type, (iii) the bump of activity is not restricted to one single SODeR cell but a group of adjacent SODeR cells is supposed to be active at the same time, (iv) the number of cells in the attractor is arbitrary in the figure and is chosen solely to illustrate the logic of wiring.

Given the huge explanatory power of the model by Navratilova et al., and the unrealistic anatomical constraints onto the MECII that an in situ, topologically 2-dimensional wiring of continuous attractor networks would require, we propose that the fundamental unit of the MECII stellate cell ensembles is a 1-dimensional ring attractor. The only substantial difference between the model and the actual wiring in the brain is the number of the neurons in the attractor. Even if we assume that the directly observed stellate cells (Gu et al., 2018) constitute just a fraction of the neurons wired together in the ring, the realistic number should be around 50 instead of 100. The correlation of the anatomical position with the spatial phase (phase of the grid) and has already been established (Gu et al., 2018). One of the strong experimental predictions of our model is that the anatomical position also correlates with the fine, theta-level temporal code. Furthermore, when the calcium imaging techniques will be perfectionized sufficiently for measurements at the appropriate time-scale and will be capable of resolving spikes in a burst, the model’s predictions regarding the in situ generation of phase precession in MECII ring attractors will be verifiable. Finally, we believe that the possibility of reactivating the bump by the ADP of the stellate cells in the position that was active in the previous theta cycle has outstanding implications for the detection of whether an event represented by the whole ring attractor (see below) has already occurred or not. Expected events, and those with a representation activated internally by a chain of already established associations can be kept continuously active this way at a late phase of the theta cycle for a longer time period, in absence of the input from other cell types such as conjunctive cells.

### 2.2. Rodent grid cells: is their spatial activity just a facette of a more general function that every reelin-positive entorhinal neuron in the layer II fulfills?

A key element of our hypothesis is the extension of the function that the reelin-positive entorhinal cells fulfill within the 1-dimensional ring attractor to the creation of new episodic memories. Such a more general function could efficiently be subserved by keeping the representations of expected events active and change their theta phase right at the time when they occur. This would relax constraints on how close in time different elements of an episodic memory have to be so that they can be joined together. Besides, we assume that there are further neurons in the entorhinal cortex that function similarly to head-direction cells but could represent any piece of relevant information and ultimately drive the activity in the ring attractors forward. The information represented by these cells is supposed to be of any complexity so that the episodic memory can the most efficiently represent what the organism encounters and perceives. In other words this would facilitate the unlimited exploitation of previous knowledge when building new episodic memories. This hypothesis is underscored by the fact that the entorhinal cortex receives highly integrated information from other neocortical areas. Further strong support to the functional extension that we propose comes from the observation that reelin-positive fan cells in the lateral entorhinal (LEC) cortex are critical to episodic-like memories (Vandrey et al., 2020). Fan cells form a network that is comparable to that of stellate cells in the medial entorhinal cortex (Nilssen et al., 2018) and their electrophysiological properties, as wells as their propensity to intrinsic oscillations are very similar to stellate cells (Canto et al., 2011). Assigning a key role to fan cells in human episodic memory would not be an assumption without a solid foundation considering that the lateral entorhinal cortex is strongly affected in preclinical Alzheimer’s disease (Khan et al., 2014). Moreover, in humans, the activity of the LEC (but not that of the MEC) is coupled to retaining the precise minute-scale temporal structure of the declarative memories (Montchal et al., 2019) and this is exactly the kind of capacity that is based on organizing experienced episodes into sequences and ultimately on the phase precession generated by the EPISODE module in our model. Nonetheless, medial entorhinal neurons can also be cue-sensitive while not displaying hexagonally arranged firing fields in the open filed (Casali et al., 2019) hence the MEC might contribute to registering episodic memories to some extent.

### 2.3. Presentation of the proposed EPISODE module

Departing from the neurons in the model of the 1-dimensional ring attractor (Navratilova et al., 2012), we propose a new entorhinal cortical module, termed the EPISODE module, responsible for the generation of the episodic memories and consisting of the following three major functionally defined cell types (figure 2):

1. *<u>E</u>vent-<u>r</u>elated <u>d</u>river* (<u>**E**</u>ReD) cell. It is an excitatory neuron in the entorhinal cortex that signals that an event is ongoing (for instance an odor is presented, an object is encountered or the head is held in a particular direction) and provides input to the PiPhA neurons. We propose that the head-direction cells, and especially those in layer III, are archetypical EReD cells.
2. *<u>Pil</u>ot of <u>ph</u>ase <u>a</u>dvancing* (<u>**Pi**</u>PhA) cell. It is an excitatory entorhinal cell that receives input both from the EReD cells and from its corresponding SODeR cell (see below). Its output excites the subsequent SODeR cells in the ring attractor, thereby moves the center of activity („bump”) in a defined direction within the ring of SODeR cells. A conjunctive grid cell in the MECIII responding to the direction of the head and displacing the bump in a ring attractor of grid cells (as described in Navratilova et al., 2012) is an archetypical PiPhA cell.
3. *<u>S</u>ignificant <u>o</u>ccurrence <u>de</u>tector <u>r</u>ing* (<u>**SODeR**</u>) cell. It is a reelin-positive excitatory entorhinal cell (stellate cell in the MECII or a fan cell in the LECII) that builds up a 1-dimensional attractor network arranged in ring topology together with other SODeR cells. Each SODeR cell fires at a defined theta phase and the phase of the activity is also changed when the excitatory PiPhA cell input arrives and changes the position of the bump within the ring attractor. A grid cell is an archetypical SODeR cell, although the observed 2-dimensional spatial activity of rodent grid cells requires additional wiring between multiple medial entorhinal EPISODE modules (see below).

### 2.4. The phenomenon of phase precession also generalized to all episodic memory items

What considerably simplifies our model and endows it with additional credibility is the fact that the generation of the phase precession emerges as its inherent property resulting from the features of 1-dimensional ring attractors the same way as in the model by Navratilova et al. (2012). We need to emphasize that there are two independent sources of phase precession: (i) the ADP of the SODeR cells is not in full synchrony with the frequency of theta, but comes somewhat faster, (ii) the PiPhA cells can displace the activity in the ring of SODeR cells when the event represented is occurring, consequently, a given SODeR cell will fire earlier in the theta cycle and thus phase precess. This explains well the bimodality of the phase precession and its steeper slope in the second part of the theta cycle (Yamaguchi et al., 2002).

Our model relies on a general EPISODE module, which is presumably ubiquitously present in all mammalian species. Therefore not only spatial representations have to phase precess, but all the representations that code for an item that can become a part of an episodic memory trace. Remarkably, earlier experiments have shown that this is exactly what happens, moreover, such phase precession has several characteristics that support our model. Recordings from rats associating a particular sound (chosen from two possibilities) with a subsequent particular odor (chosen from two possibilities) and then completing a task have accordingly demonstrated that CA1 neurons representing an odor start to phase precess when the odor is presented (Terada et al., 2017). Furthermore, choice selective cells start to phase precess when the choice is actually made based on the particular odor-sound combination. Importantly, choice selective cells start to fire at 270° when the second cue is presented, and phase precess down until 90° when the time window for the choice opens. On the other hand, odor-selective cells start to fire at 270°, when the first cue (sound) is already presented and the odor can be expected. Odor-selective cells are re-set to 270° when the choice is completed, and, interestingly, the figures from the study seem to show (although the authors do not state this explicitly in the text), that this is achieved abruptly but still in a „circular” way: spikes are first placed to a phase earlier than 90°, and then to a phase slightly higher than 270° (Terada et al., 2017). These observations are all consistent with an entorhinal 1-dimensional ring attractor generating phase precession and points to the 270° being the phase of anticipation. In addition, an early report (Pastalkova et al., 2008) of what later were termed CA1 time-cells (MacDonald et al., 2011) described that these neurons also phase precess within their temporal window of non-spatial activity, and the slope of phase precession is inversely proportional to the length of their activity window (Pastalkova et al., 2008). This suggests that phase precession of approximately 180° is a fundamental unit that corresponds to the occurrence of an event and the slope of the phase precession can be adjusted according to how long the event takes. Taken together, these observations strengthen the already formulated hypothesis that phase precession is the neural code underlying episodic memories (Jaramillo and Kempter, 2017) and other brain areas may inherit phase precession from the entorhino-hippocampal system. This view is fully consistent with our model and is mechanistically explained by the functioning of the EPISODE module (see above) and the downstream processing in the hippocampus proper (detailed below).

An especially remarkable feature of scopolamine, from the point of view of the model presented here is that it almost completely suppresses phase precession in the CA1 region of the hippocampus without changing the spatial activity of individual cells, nor eliminating the theta rhythm. The effect is detectable upon a single pass through a place field of a single CA1 neuron (Newman et al., 2017). The other, complementary piece of the puzzle is that scopolamine causes very serious anterograde amnesia in human subjects with its effects being stronger than those of diazepam (Ghoneim and Mewaldt, 1975) and correspondingly is also used by criminals (Reichert et al., 2017). These two correlating effects of scopolamine provides a further strong argument that phase precession could be indispensable for creating episodic memory traces.

The function of the entorhino-hippocampal system is not limited to remembering: intelligent planning and pondering the outcome of various future alternatives can also rely on this structure (Kay et al., 2020; Vikbladh et al., 2019). Locking at 270° those phase precessing elements that are expected may assist this additional function. As observed (Pastalkova et al., 2008; Terada et al., 2017), this kind of locking only happens when the particular event is truly expected to happen and is built into a task meaningful for the animal. Ctip2 positive neurons from the layer Vb of the entorhinal cortex constitute a candidate cell type to mediate such expectation, since they project to the stellate cells in the MECII, and receive input from the CA1, subiculum and retrosplenial cortex.

### 2.5. Generalization from grid cells to SODeR cells

One key element of our model is the phase code generated by entorhinal 1-dimensional ring attractors and used by the downstream hippocampal DG-CA3-CA1 network to link together elements with different phases. An obvious requirement is that these elements have to represent many different kinds of, in many cases non-spatial, events of an episodic memory. Some SODeR cells have to represent objects, sounds, odors, places, or even composite events already linked together at the neocortical level, and, on the other hand, the theta phase assigned to a particular event should clearly reflect whether it is in the future (expected), present or past. Rodent experiments permit the relatively easy identification of object-specific neurons in the lateral entorhinal cortex (Tsao et al., 2013), furthermore, higher-order properties of objects not directly derived from available stimuli are represented by these cells such as the previous presence of a particular object during a particular trial in the past (Tsao et al., 2013). These observations make lateral entorhinal fan cells ideal candidates for SODeR cells required for episodic memories. Further experiments are needed to clarify the contribution of other SODeR cells to episodic memories, since neurons have been identified in the medial entorhinal cortex that are responsive to cues and do not display firing fields with six-fold symmetry at all in the open field (Casali et al., 2019). More specifically, cue cells were identified both in the MECII and in the MECIII (Kinkhabwala et al., 2018), moreover, they were arranged in clusters (Kinkhabwala et al., 2018) similarly to grid cells (Gu et al., 2018).

### 2.6. Generalization from head direction cells to EReD cells

Our model postulates that the head-direction cells providing input to the conventional grid cells have to be sensitive to the direction of the movement and its speed. In harmony with this assumption, head-direction cells with firing rate proportional to the speed of the animal have been described (Hinman et al., 2016). In the MECII, the population of pyramidal cells, located in the calbindin positive patches, show considerably stronger speed modulation than the stellate cells in the same layer, and the fraction of the cells responsive to the speed of the animal is much larger in the pyramidal patches (Sun et al., 2015). Therefore a fraction of calbindin positive pyramidal neurons could also be EReD cells, or could provide input to EReD neurons located elsewhere in the medial entorhinal cortex.

EReD cells in the LEC are postulated to fire action potential bursts upon the presentation of their congruent object based on the assumed analogy with head direction cells in the MEC. Further experiments are needed to tell them apart from, and identify their connections with local cue-sensitive circuits.

Our model can provide a potential explanation for an intriguing previous finding related to head-direction cells. It is reasonable to assume that the bump within the 1-dimensional ring attractor of the grid cells can travel in the two opposite directions. To better represent the position of the animal, deteriorative noise could be filtered out by segregating PiPhA input for the leftward and for the rightward movement along a given axis of the hexagonal grid. This kind of segregation could avoid the simultaneous activity of two different PiPhA cell population driving the bump in two opposite directions (figure1). Mechanistically, this would necessitate two antagonistic populations of EReD cells similar to the „north” and „south” head-direction cells in the model by Navratilova et al. (2012). Strikingly, this could very well be achieved by the experimentally found theta-skipping properties of the head-direction cells in rats foraging in open fields (Brandon et al., 2013). In that study, some head-direction cells were found to skip a theta cycle completely, and populations of head-direction cells firing on alternate theta cycles were systematically found to code for different directions, although the angle between the two directions was not equal to exactly 180° (Brandon et al., 2013). Still, theta-phase skipping segregates neurons into two subpopulations with mutually exclusive activity, one subpopulation being active on even, and the other on odd theta cycles.

### 2.7. Generalization from conjunctive cells to PiPhA cells

Conjunctive grid cells, in other words neurons that are sensitive to head-direction and, at the same time, display firing fields arranged into a hexagonal grid, can routinely be detected in the medial entorhinal cortex (Sargolini, 2006). Conjunctive grid cells are typical PiPhA cells that are arranged in a topology that follows the 1-dimensional ring attractor of the SODeR cells but do not excite each other. We predict the presence of non-spatial cue-sensitive PiPhA cells in the superficial layers of LEC.

Theta phase skipping must be observed also among PiPhA cells if each 1-dimensional ring attractor is driven by two antagonistic sets of them, in line with our assumption presented above regarding the EReD cells. Indeed, recordings from the MEC of rats have shown that conjunctive grid cells display theta phase skipping (Hinman et al., 2016). In the same report, omnidirectional grid cells do not seem to skip theta cycles (figure 3), although this could only be deducted from the figure and the overall number of theta modulated cells reported (Hinman et al., 2016).

**Figure 3.**
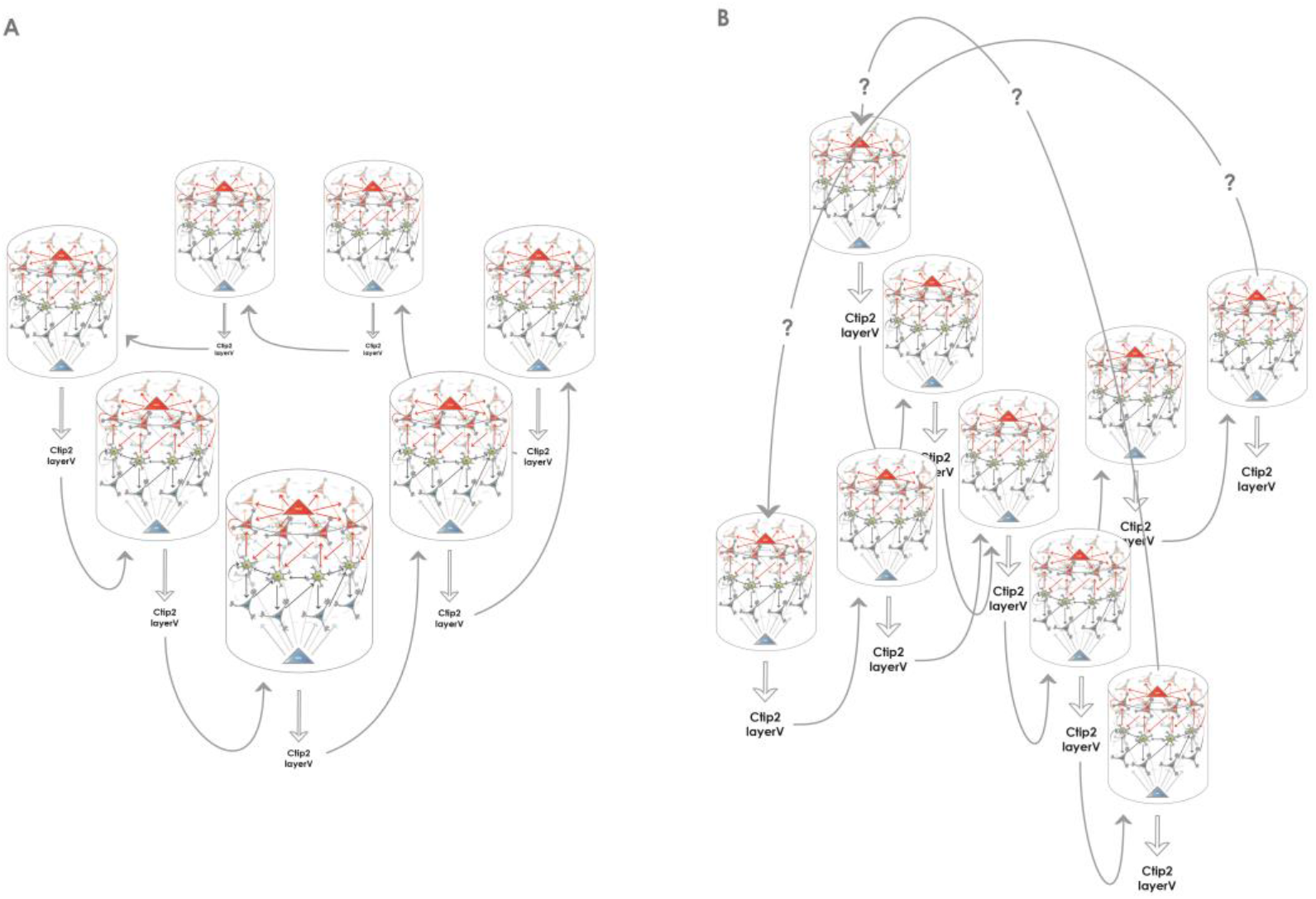
Creating higher dimensional units from multiple 1-dimensional ring attractors. **A**: Medial entorhinal 1-dimensional ring attractors could collaborate to form a 2-dimensional network that accounts for rodent grid cell activity. Each 1-dimensional ring attractor sends information to the layer V of the medial entorhinal cortex to neurons such as Ctip2 positive pyramidal cells that relay this information to a neighboring 1-dimensional ring attractor. The structure could encircle the calbindin positive pyramidal patches, however, this kind of arrangement is not required for the model to be functional, the wrap-around mechanism for the extra dimension is still assured. **B**: Medial entorhinal 1-dimensional ring attractors could collaborate to form a 3-dimensional network that can better subserve the hexagonal symmetry of the firing fields that rodent grid cells display. The communication between the 1-dimensional units is similarly possible via the layer V, however, the way the wrap-around connectivity is implemented is less straightforward in this case.

One single piece of evidence points to the fact that ring attractors may generally be driven in two directions, that is, bidirectionality would not be restricted to ring attractors composed of bona fide grid cells. In an ingenious experiment, rats were trained to run forward on a treadmill that was mounted on a miniature train transporting them either forwards or backward along a track (Cei et al., 2014). Strikingly, not only theta sequences displayed by CA1 cells were reversed during backward transport but also phase precession plots (ie. circular-linear correlation plots) had opposite slopes (Cei et al., 2014). We will shortly describe in chapter 3. how the CA3 could contribute within these conditions to the reversal measured in the CA1 region.

### 2.8. How is the 2-dimensional attractor network required for rodent grid cell activity established?

We are aware that the 1-dimensional ring attractor of SODeR cells cannot account in itself for the highly regular 2-dimensional spatial activity of the grid cells (figure 1). Continuous attractor network models typically assume a 2-dimensional sheet of neurons within which the activity spreads as the rodent navigates in a 2-dimensional environment (McNaughton et al., 2006). What we propose tentatively to resolve the contradiction is the linking of 1-dimensional ring attractors together in a way that introduces a second dimension. Calcium imaging has shown that within a grid module, stellate cells functioning as grid cells are arranged into at least five neighboring phase clusters that work in synchrony to some extent (Gu et al., 2018). This structure seems to repeat in all modules and the clusters show some functional independence, since firing fields can be skipped in one cluster while being present in the neighboring one (Gu et al., 2018). Our assumption, therefore, is that when the animal moves in a direction orthogonal to the one coded by the 1-dimensional attractor, the bump is displaced to an adjacent ring of grid cells that corresponds to the neighboring cluster of active neurons observed by Gu et al. (2018). In the absence of experimental findings, only hypotheses can be formulated regarding the mechanism of such displacement, nonetheless, the communication with deeper layers of the medial entorhinal cortex might be involved. For instance Ctip2 positive cells in the layer V have reciprocal connections with superficial stellate neurons (Witter et al., 2017) and hence might receive input from one ring attractor and transmit the signal to an adjacent one. This kind of mechanism would also resolve the conundrum of wrapping around along the second dimension since the directly observable clusters are next to each other (Gu et al., 2018) and can readily form a circle (figure 3A). We are aware that for the phase precession to be nearly omnidirectional, additional mechanisms might have to be postulated. Further uncertainty comes from the fact that the hexagonal lattice of grid cell firing fields might be more efficiently created by a 3-dimensional continuous attractor network composed of juxtaposed 1-dimensional ring attractors arranged in an orthogonal lattice, although the wrapping around is anatomically more challenging in this case (figure 3B). Therefore the model presented in this paragraph is adumbrative, nonetheless, it supplies a strong and directly testable prediction, namely the systematic jumping of the activity between the clusters when the animal moves in the appropriate direction in an open field. The previous observations that the grid pattern is elliptically distorted (Stensola et al., 2015), furthermore it is compressed congruently along a particular axis (but not along the other one) when the arena is compressed (Munn et al., 2020) support the core idea of using the 1-dimensional ring attractors as fundamental building blocks and linking them to add extra dimensions. Finally, we propose that the regular cytoarchitecture of the MEC, which contrasts the anatomy of the LEC, is linked to the way that the higher dimensional ring attractors are built from 1-dimensional building blocks.

## 3. The hippocampal component of the episodic memory circuit: creating associations and organizing sequences

### 3.1. General principles of the proposed model

The entorhinal mechanisms described in the previous chapter assign a theta phase to each relevant event that is to be incorporated into an episodic memory trace. Such an event can in the first place be detected by any sensory modality, and its complexity can vary according to the extent of processing by neocortical areas before being passed on to the entorhinal cortex and acquiring the form of a phase precessing signal. Further processing is performed by the hippocampus proper that creates a stable association between the events and organizes them into a meaningful sequence that can be read out from the hippocampal CA1 in the form of a theta sequence during wakefulness and as a more compressed ripple-associated sequence during slow-wave sleep. Several propositions in the literature deal with the details of the associative and sequence organizing mechanisms in the hippocampus, below we only describe these mechanisms to the extent needed to build a coherent model of registering new episodic memories.

### 3.2. The entorhinal cortex → dentate gyrus processing: pattern completion and pattern separation

The information from the SODeR cells of the entorhinal cortex is sent to the granule cells of the dentate gyrus (DG), where pattern separation and, in some cases, pattern completion is achieved (Nakashiba et al., 2012). Relatively recent findings have clarified that young granule cells mediate pattern separation while new granule cells, generated shortly before the time of registering the new episodic memory, can mediate pattern completion (Nakashiba et al., 2012). This view is fully consistent with our core hypothesis according to which information from the SODeR cells is directly passed on to granule cells, and the newborn ones of the latter are receptive to form new synapses from ring attractors that they are not stably connected with yet (Piatti et al., 2013) but are active at the time of the occurrence of the event to be memorized. The relatively short time window during which the newborn granule cells are receptive lays the foundation of the pattern separation based on the wiring between the entorhinal cortex and a particular young granule cell (figure 4). On the other hand, old granule cells can only use their already established contacts, therefore an episodic memory can be recalled when all of, or the majority of the entorhinal inputs synaptically bound to a given granule cell are reactivated. This provides a clear-cut explanation why the memory engrams from a particular temporal window can be best captured at the level of a granule cell population (Liu, 2012; Redondo et al., 2014, Ramirez et al., 2015, Cazzulino et al., 2016): the experimental reactivation of a sparsified code that strictly represents only the elements bound together at the registration of the episodic memory (and the smallest possible amount of other unrelated elements) is sufficient to reactivate that particular memory and can even influence animal’s behavior in a congruent manner (Cazzulino et al., 2016; Liu et al., 2012; Ramirez et al., 2015; Redondo et al., 2014). Furthermore, this model sheds light on why pattern completion processes are experimentally found to be mediated by old dentate granule cells (Nakashiba et al., 2012): when the set of the environmental cues is submaximal but only a small portion of them is missing (ie. submaximal number of SODeR cells send input to the granule cells, as an example, 3 out of the 4 cues in figure 4. are activated), the corresponding granule cells might still be driven over the activation threshold and this way the memory is recalled since the activation of the whole downstream CA3 network is assured through the very potent detonator synapses.

**Figure 4.**
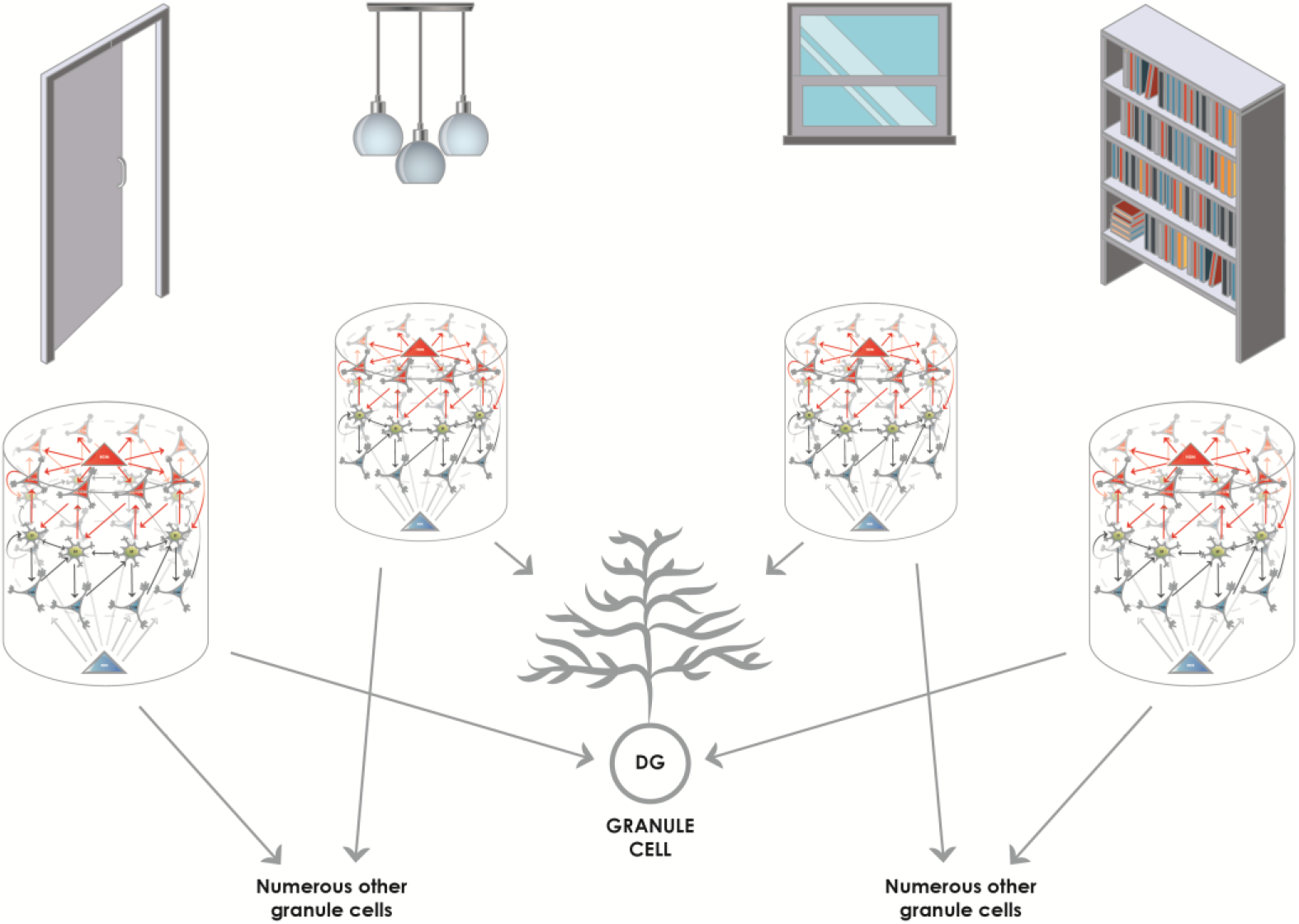
Converging and diverging routes of the information from the entorhinal cortex to the granule cells of the DG. Reelin-positive neurons (functionally SODeR cells in our model) provide excitatory input to the granule cells of the DG. Newborn granule cells are particularly receptive within a defined temporal window of their maturation to form connections with incoming fibers that are active. Therefore the elements represented by the 1-dimensional ring attractors can be associated together, and the convergence assures that granule cell activity will be highly specific to the combination of the elements encountered. Of note, the elements themselves can be of any complexity given the different levels of integration in the entorhinal cortex. It is reasonable to assume that early on elements like simple objects can be learned by associating their unique features based on this mechanism. Later such representations could be semanticized and more complex episodic memories based on already learned objects can be created the same way. The figure illustrates this latter stage of learning, and shows how objects in a room (door, pendant, window, and shelf) can be associated together to be part of an episodic memory. The results from Nakashiba et al. (2012) show that old granule cells can mediate pattern completion to some extent, this would be explained by the fact that a slightly reduced number of active elements (such as three out of the four elements represented in the figure) is still capable of activating a given granule cell that established contacts with a full set of elements (four in the figure) during its young and receptive period.

To preserve the fine, millisecond-scale temporal code generated by the LECII, specific wiring must be postulated: the same granule cell in the DG must be targeted by all of the SODeR cells in a given 1-dimensional ring attractor coding for events (sounds, odors, stimuli, cues, etc.) and this way the phase precession can be transmitted to the hippocampus (figure 5). However, this particular way of wiring is not supposed to apply to grid cells in the MECII. The assumption that all the fan cells that are part of the same cue-sensitive 1-dimensional ring attractor target the same granule cell constitutes an important and testable prediction of our model. Of note, this way only events having similar theta phases could be linked together efficiently at the level of granule cells, according to the scheme in figure 4. Therefore we propose a division of labor for establishing associations and completing patterns: the DG performs these operations for strictly coincident events while – as detailed below – the CA3 for sequential ones.

**Figure 5.**
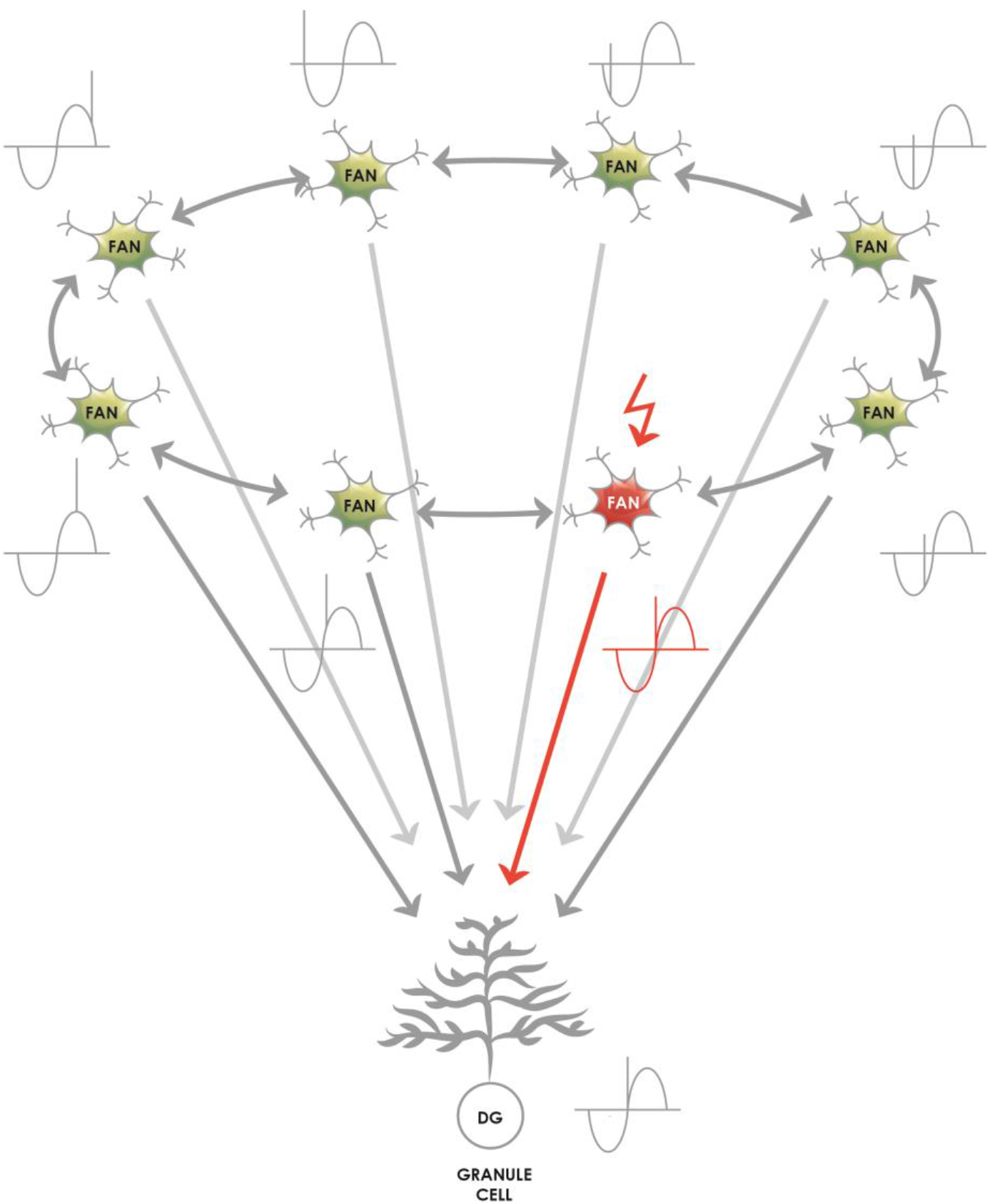
The proposed mechanism by which the theta phase code generated by the lateral entorhinal 1-dimensional ring attractors is passed on to the granule cells of the DG. The 1-dimensional ring attractors are naturally capable of generating a signal that shows theta phase precession at the level of their individual neurons (Navratilova et al., 2012) and phase precession originating in the ECII is passed on to the hippocampus (Schlesiger et al., 2015). The representation of relevant events (such as cues, odors, sounds, places and complex events in rodent experiments) shows phase precession, and a reasonable assumption is that a lateral entorhinal 1-dimensional ring attractor consisting of many SODeR cells represents an event and generates the event-specific phase precessing signal. As a consequence, the target granule cell must be informed of the current theta phase equally during the anticipation, the onset and the termination of the event. The most likely and simplest network wiring compatible with this requirement is an excitatory connection from each fan cell of a given 1-dimensional ring attractor to a single granule cell. The figure shows that the theta phase of the currently active SODeR cell (fan cell in red) determines the theta phase at which the target granule cell fires. We would like to emphasize that such wiring is neither required nor plausible for rodent grid cells despite their similar organization into 1-dimensional entorhinal ring attractors.

### 3.3. The DG → CA3 and CA3 → CA3 processing: sequence generation and sequence-based pattern completion

From the point of our hypothesis, the information processing in the recurrent collateral system of the CA3 region is of paramount importance. The fine temporal code generated by the ring attractors of the SODeR cells can be read out by the CA3 via the following mechanism: spikes at a given theta phase could appear on the recurrent collaterals while spikes at neighboring theta phase could appear at the same time as mossy fiber input. At least the physiology of the CA3 neurons and their apical dendrites permits the turning of subsequent inputs into coincident ones this way (Raus Balind et al., 2019). The input coming in at earlier theta phases is assumed to correspond to events that have already been moved forward in the ring attractor of SODeR cells by PiPhA cell activity and thus precede ongoing or expected events. After running through the recurrent collaterals, such input will be coincident at the CA3-CA3 synapses with a second input representing ongoing events and arriving at a later theta phases (figure 6A). Therefore the CA3 circuit is perfectly positioned to forge sequences from spikes (events) that have subsequent theta phases, and also to bind those events together via the NMDA-dependent plasticity at the synapses between CA3 recurrent collaterals and CA3 apical dendrites (figure 6A). Ultimately „theta sequences” are constructed this way by reinforcing the connection between each subsequent pair of relevant events.

**Figure 6.**
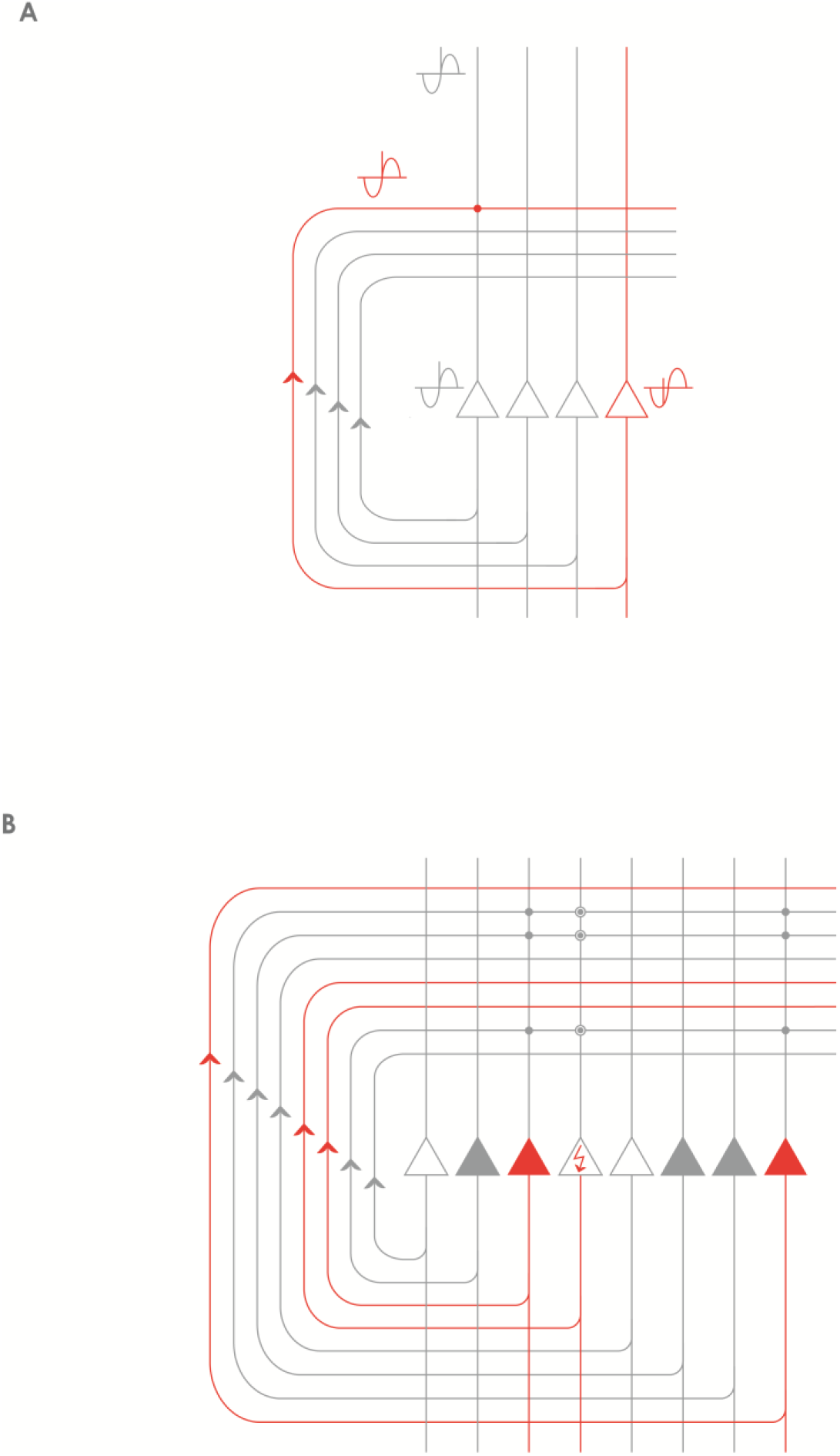
Processing of the phase precessing signals by the recurrent CA3 network. **A**: Signals that are subsequent along a theta cycle but arrive within a defined window of time can be made coincident by the recurrent collateral CA3 network. The event represented by the red CA3 neuron (and the remaining unshown neurons of the corresponding population) is previous to the event represented by the grey CA3 neuron on the left (and the remaining unshown neurons of the corresponding population) as evidenced by its earlier theta phase. However, the time it takes for the action potentials to travel along the CA3 recurrent collateral axons can make the red and the grey representations coincident at the level of CA3-CA3 synapses. **B**: The recurrent CA3 network is especially well-positioned to perform pattern completion: the event represented by the grey CA3 neurons can complete the representation of the event the red CA3 neurons code for. The neuron depicted with the red arrow in the perisomatic region is not activated by the external cues but it can still be activated by the population of the grey neurons via the synapses potentiated by previous experience (encircled grey dots). Best pattern completion is assumed to be achieved in the CA3 on one hand for events already organized in a sequence and on the other hand by an event that is previous to the event with incomplete representation.

Of note, an outstanding publication provided proof that in the CA1 region, at the very first encounter with a linear track, single-trial phase precession always occurs, and seems to be an intrinsic property of the circuitry, while organized theta sequences appear only at the second run (Feng et al., 2015). Later a further publication added a firm piece of evidence that dissociation between phase precession and theta sequences is possible (Middleton and McHugh, 2016) and provided proof that the emergence of organized theta sequences in the CA1 is dependent on the CA3 input (Middleton and McHugh, 2016). That observation is fully compatible with our model that locates the source of the hippocampal phase precession in the entorhinal ring attractors of SODeR cells and such phase precession is assumed to be independent of experience, while, on the other hand, organized theta sequences are proposed here to arise in the CA3, although might be projected back to the EC via the CA1 region. It is not clear what phase difference is needed for two events to be recorded as subsequent elements of a sequence. As proposed originally, gamma oscillation could define at what pace a new subset of CA3 neurons could become active (Lisman and Jensen, 2013), therefore the actual frequency of local gamma oscillation might determine how separated two events have to be along the theta cycle to be registered by the CA3 as two subsequent events of a theta sequence.

Using rodent experiments, a pioneering publication reported that plasticity at the CA3-CA3 synapses is required for heteroassociation (binding different cues together) in the Morris Water Maze task (Nakazawa, 2002). However this kind of learning might also involve some sequences provided that the animal discovers the cues in a sequential manner. Indeed, later it has been shown that learning sequences of odors presented at a 3 seconds interval relies exclusively on CA3, and does not rely on CA1 (Farovik et al., 2010). In line with this finding the major function that we propose here for the CA3 network is the memorizing of new sequences.

Traditionally the pattern completion needed for the retrieval of an episodic memory based on an incomplete set of cues is thought to be carried out by the CA3 region, and population code read out from the DG and the CA3 confirmed this long-standing view (Neunuebel and Knierim, 2014). What we suggest here is that the CA3 can participate in the pattern completion provided that the elements of the memory were previously organized in a temporal sequence. In this case, earlier elements of the sequence can not only evoke the representation of a subsequent element, but also complete it by activating the CA3 neurons missing their appropriate input (figure 6B). It has to be noted though, that, as detailed above, the DG is also shown to contribute to the pattern completion process.

A remarkable property of the STDP at the recurrent CA3 → CA3 synapses is the almost complete simmetricity, in other words the lack of strong preference for the temporal order of the presynaptic and postsynaptic input (Mishra et al., 2016) which differs from STDP elsewhere in the brain. Thus a discrete attractor state of the CA3 network representing a snapshot-like element of an episodic memory could theoretically activate either the previous or the subsequent attractor state. This provides a potential explanation for the fact that in the brains of rats transported backward along a track, not only the slope of the phase precession was inverted, but also the CA1 theta sequences unfolded in the opposite order (Cei et al., 2014).

The sequences of events that are written into the local CA3 circuit by the mechanism described above are likely to be those that are replayed in sharp-wave ripple (SWR) events during waking immobility (eSWR) and in slow-wave sleep (sSWR) and trigger downstream CA1 sequences. This is in harmony with the fact that SWRs were early on proposed to be generated in the CA3 region (Csicsvari et al., 2000), and later in vitro experiments confirmed this proposition (Schlingloff et al., 2014). In vivo rodent experiments have shown that spatial positions represented by CA1 ensembles reactivated during the eSWR events are discretely distributed along the whole trajectory that the reactivation codes for, and, most importantly, both the step size and the frequency of jumps seem to be governed by CA3 attractor dynamics in a way that one particular position seems to correspond to one particular discrete attractor state of the CA3 network (Pfeiffer, Brad, 2012). Finally, reactivating sequences of CA1 neurons during eSWR events can code for a trajectory that is the opposite of the one actually taken by the animal. These CA1 sequences are termed reverse replays (Ambrose et al., 2016; Foster and Wilson, 2006) and are likely to be the manifestation of the bidirectional link between the neighboring discrete CA3 attractor states. Taken together, sequences written into the recurrent CA3 circuit are the most likely source of the information replayed in sSWR events that broadcast information to neocortical areas, especially to the so-called default mode network, and promote the consolidation of recently acquired memories in sleep (Walker and Robertson, 2016).

### 3.4. The CA3 → CA1 processing: integration, comparison and direct retrieval

Input is mainly provided to the CA1 area through two major pathways that target distinct regions of the CA1 dendrites. First, the Schaffer collaterals convey information from the CA3, which is already temporally organized at the theta timescale and patterns are already processed via the pattern separation and pattern completion activity of the DG. Second, the temporoammonic pathway directly originating from the ECIII provides information that underwent different depths of integration nonetheless represents the current external and internal state of the organism. Our model postulates that the major role of the CA1 is to associate the particular pieces of information, arriving via the CA3 network, into a broader context. This might be achieved via linking co-active CA3 neurons to a common CA1 target given the substantial CA3 → CA1 convergence, but more straightforwardly, the actual context could also be represented by the temporoammonic pathway.

In line with this idea, the direct integration of the ECIII and CA3 input has experimentally been verified by measuring the so-called plateau potentials in the CA1 cells (Bittner et al., 2015). Of note, a gating mechanism in the CA1 can emphasize either one or the other of the two inputs (Leão et al., 2012). What is important for the proposed model, is that the CA1 neurons receiving input from the medial entorhinal cortex are highly context-sensitive even though they fire at specific landmarks in a given context (Geiller et al., 2017).

Besides the integrative function described above, the CA1 can compare sequences stored in the CA3 with the actual state of the environment. This is probably best exemplified by the fact that the CA1 neurons code for non-spatial sequences such as odor sequences and can detect when a mismatch comes in an already learned sequence (Allen et al., 2016). A strikingly high proportion (close to 25%) of the CA1 neurons show differential activity upon the presentation of the correct or of a non-matching element of a memorized odor sequence (Allen et al., 2016). This finding inspired a model (Barrientos and Tiznado, 2016) where the learned sequence is replayed by the trisynaptic loop, while the actual element presented is represented by the temporoammonic pathway and the CA1 decides about the correct or incorrect nature of that particular element.

A similar function of the CA1 is likely to contribute to evaluating future options. Experimental evidence suggest that CA1 sequences detected within SWR events can predict the path that rodents take to navigate to learned goals (Pfeiffer and Foster, 2013), furthermore, can also represent a path that the animal has never actually taken (Gupta et al., 2010). It is thus reasonable to speculate that the sequences arising in CA3 are not only used by the CA1 for the purpose of comparison with past events, but also for intelligent planning. This hypothesis is supported by some initial experimental findings showing that the hippocampus contributes to model-based planning (Vikbladh et al., 2019). We, therefore, propose that the rigidity by which one discrete attractor state of the CA3 triggers the next one is reduced and there is at least some level of flexibility of jumping to an alternative subsequent discrete CA3 attractor state.

As compared to other regions of the hippocampus proper, the connectivity of the CA1 is unique in the sense that it receives direct input from the ECIII. It has been shown that fast gamma oscillations contribute to the transmission of information from the ECIII while slow gamma from the CA3 (Colgin et al., 2009). The prevailing view is that currently available information, provided by internal and external stimuli, reach the CA1 from the ECIII while recalled information arrives from the CA3. Nonetheless our model is also compatible with a view that CA1 can retrieve non-sequential information by relying solely on the ECIII input. This kind of direct retrieval would be dependent on the prior reinforcement of the temporoammonic input on a CA1 cell by the coincident inputs from different EPISODE modules via the trisynaptic loop; and the prior, concomitant reinforcement of the connections such CA1 cell has with its downstream targets.

## 4. Summary and conclusions

The human medial temporal lobe system has the remarkable capacity of storing experiences encountered only once and this observation can most likely be generalized to the majority of the mammalian species since one-time learning is documented in experimental animals as well. Our model describes how elements of new episodic memories are associated together and how pattern separation and pattern completion is achieved during this process. We describe a new functional unit in the entorhinal cortex that we term the EPISODE module. We define it as a 1-dimensional ring attractor network composed of reelin positive neurons in layer II and of two further cell types that cooperate to displace the activity within the ring. Together, the three cell types are proposed to create a fine, millisecond scale temporal code that, in turn, enables the organization of memory elements into a meaningful sequence in the CA3. The power of the EPISODE module lies in explaining on the one hand side the emergence of the strikingly regular grid cell activity in rodents and on the other hand the generation of the phase precession that propagates from the entorhinal cortex through all of the neuronal populations of the hippocampal trisynaptic loop. We cite a large amount of experimental data that supports the generalization of the way that all phase precessing elements of any episodic memory are organized into sequences in the hippocampus. Importantly our model supplies a set of experimentally testable predictions that are listed in table 1.

**Table 1.** Experimentally testable hypotheses derived from the model.

| **<u>Hypothesis</u>** | Reference with experiments similar to the proof-of-concept experiment | Required improvement of the currently available experimental techniques |
| --- | --- | --- |
| The anatomical position of the SoDeR cells correlates with the theta phase of the spikes they emit (similarly to correlations between the anatomical position and the spatial phase in case of grid cells). | Gu et al., 2018 | Calcium imaging with finer temporal resolution |
| All of the SoDeR cells in a particular cue-sensitive 1-dimensional ring attractor of the LECII target the same granule cell in the dentate gyrus. | - | Retrograde labelling based on optogenetic tagging and activation of a DG engram |
| Systematic jumping of the activity between grid cell clusters of the same grid cell module in the MECII in moving animals (the presence of a 2 or 3-dimensional continuous attractor neuronal network). | Gu et al., 2018 | - |
| The LECII contains cue-sensitive ring attractors, within which the activity is strongly displaced when the congruent cue is presented (odor based experiments coupled to imaging would be ideal to test this hypothesis). | Terada et al., 2017<br>Gu et al., 2018 | - |
| Conjunctive grid cells show theta phase skipping and pairs of them active on alternate theta cycles are responsive to different directions. | Brandon et al., 2013 | - |
| During backward transport of rats running forward on a treadmill, the activity propagates backwards within cue-sensitive ring attractors in the LECII | Cei et al., 2014<br>Gu et al., 2018 | Calcium imaging compatible with forced transport while treadmill running |

## Acknowledgment

Hereby I would like to express my thankfulness to Dr. Jozsef Csicsvári, Dr. Joseph O’Neill and Dr. Philipp Schönenberger for the fruitful coffee discussions during my previous research position at the Institute of Science and Technology (Austria) that led to great ideas which ultimately served as the groundwork for some of the concepts presented in this article.

